# An integrated RNA-proteomic landscape of drug induced senescence in a cancer cell line

**DOI:** 10.1101/2023.03.21.533605

**Authors:** Thomas Stevenson, Maha Al-Roshdi, Franziska Görtler, Sushma Nagaraja Grellscheid

**Author notes:** these authors contributed equally to this work (Thomas Stevenson, Maha Al-Roshdi). these authors share senior authorship (Franziska Görtler, Sushma Nagaraja Grellscheid).

## Abstract

Senescent cells are characterized by an arrest in proliferation. In addition to replicative senescence resulting from telomere exhaustion, sub-lethal genotoxic stress resulting from DNA damage, oncogene activation, mitochondrial dysfunction or reactive metabolites also elicits a senescence phenotype. Senescence is a controlled programme affecting a wide variety of biological processes with some core hallmarks of senescence as well as tissue specific changes. This study presents an integrative multi-omic analysis of proteomic and RNA-seq from proliferating and senescent osteosarcoma cells. This study demonstrates senescence induction in a widely used cell line which can be used as a model system for characterising cancer cell responses to sub-lethal doses of chemotherapeutic agents, and makes available both RNA-seq and proteomic data from proliferating and senescent cells in open access repositories to aid reuse by the community.

## Introduction

Cellular senescence describes the irreversible loss of cells’ ability to proliferate. This phenomenon was first described by Hayflick and colleagues six decades ago when they showed that cultured human fibroblast cells have a limited capacity to proliferate [1]. This durable cell cycle arrest is resistant to mitogenic stimuli and distinct from other hyporeplicative states such as quiescence or terminal differentiation. First considered an artefact of in vitro cell culture, senescence is now considered a fundamental cellular process associated with a broad range of developmental and pathological processes and an irrefutable hallmark of organismal ageing [2, 3].

Replicative senescence is induced through telomere attrition, however senescence may also be induced by several other intrinsic stressors, such as oxidative stress, oncogene activation, or genomic instability [4]. Senescence is also induced through extrinsic stimuli, including viral infection, radiation, and chemotherapeutics [4, 5]. The onset of senescence is associated with a range of molecular and morphological traits, including the expression of several senescence markers and a significantly enlarged and flattened appearance [6, 7]. The most commonly employed method to identify senescent cells is to stain for senescence-associated ß-galactosidase activity (SA-ß-gal). This assay exploits the unique pH of senescent cell lysosomes (pH 6.0), which can be detected using X-gal staining. Additional molecular characteristics include the formation of senescence-associated heterochromatin foci (SAHF) and the activation of p53 and p21, occurring at the onset of cell cycle arrest, which is subsequently maintained by the constitutive activation of p16 [8, 9].

Cellular senescence is classically considered an anti-tumour mechanism, acting as a barrier to proliferation in the event of significant cellular damage [10]. Although unable to divide, senescent cells remain metabolically active, allowing them to participate in a range of physiological processes and secrete a range of potent inflammatory proteins known as the senescence-associated secretory phenotype (SASP) [11]. The SASP is thought to have evolved to aid in eliminating senescent/neoplastic populations through the recruitment of immune cells [3, 12]. However, as the number of senescent cells increases with age, the constant production of these inflammatory proteins promotes tissue dysfunction and contributes to a range of age-related diseases such as cardiovascular disease, diabetes, and cancer [11, 13, 14, 15]. Thus, senescence is highly pleiotropic and exists as part of a complex balance to maintain the function and health of cells, tissues, and organisms.

Senescence and apoptosis are closely linked and together form the primary protective mechanisms to suppress tumorigenic events. Dysregulation in the apoptotic apparatus is well described during neoplastic transformation, and there is increasing evidence that pathways of senescence induction are also inhibited [16]. Several chemotherapeutic drugs have been shown to induce senescence in cancer cells. An example is the topoisomerase inhibitor doxorubicin, used clinically to treat cancers of the blood, stomach, lungs, and ovaries (amongst others), which disrupts the re-ligation of DNA strands and leads to the activation of the DNA-damage response [17, 18, 19]. Although inducing a non-proliferate state in tumour cells may appear to be a favourable outcome, it is likely to be a heterogeneous response with one population entering complete senescence whilst others continue to proliferate [16]. Moreover, the induction of senescence and subsequent secretome produce a microenvironment conducive to tumourigenesis and/or disease relapse [16]. It is also possible that this heterogeny gives rise to a more aggressive tumour population.

Understanding the mechanisms through which cancer cells can escape proliferative arrest which is otherwise definitive in surrounding normative cells is essential to understanding how malignant cells resist genotoxic drug therapy [16]. Here we have integrated RNA-Seq and proteomic analyses to investigate the transcriptome of a model of chemotherapy-induced senescence in a common cancer cell line. We have induced senescence using doxorubicin, tracked the development of senescence and compared our data to existing data in commonly used models of cellular senescence.

## Methods

### Cell Culture

Cells were maintained at 37°C, 5.0% (v/v) CO2, and ∼95% humidity and passaged when they reached ~90% confluency. All cells were cultured in Dulbecco’s Modified Eagle’s Medium supplemented with 10% foetal bovine serum, 2 mM L-glutamine, 100 U/ml penicillin and 100 ug/ml streptomycin [20].

### Senescence Induction and Identification via Senescence Associated *β*-Galactosidase Detection

Senescence was induced via incubation in 200 nM doxorubicin for 48 hours. The media was then exchanged and the cells cultured for an additional 5-7 days. Senescent cells were identified via staining with SA-*β*-Gal staining solution (150 mM NaCl, 200 mM MgCl2, 40 mM citric acid, 12 mM sodium phosphate, 5 mM potassium ferrocyanide and 5 mM potassium ferricyanide, adjusted to Ph 6.4). Cells were fixed in 4 percent PFA prior to incubation with the staining solution overnight at 37°C. Cells were imaged using a bright field Evos XL Core Cell Imaging microscope.

### SWATH-MC

Cells were washed in PBS and lysed in RIPA buffer (150 mM NaCl, 1pc Nonidot P-40, 0.1 pc SDS, 0.1 pc sodium deoxycholate, 50 mM Tris (pH 7.4)) and centrifuged at 20,000 × g for 20 min at 4°C. The protein concentration of the supernatent was measured using a DC protein assay allowing the manufacturer’s instructions.

Preparation of peptide samples for proteomic analysis and mass spectrometry was performed by a department facility. Briefly, samples were prepared using FASP Protein Digestion Kit and sequencing grade-modified trypsin. The samples were then freeze-dried and re-suspended in 3% acetonitrile, 0.1% TFA followed by a de-salting step. Sample fractions (5 ug peptides) were analyzed using ekspert TM nanoLC 425 coupled to a quadrupole Time-Of-Flight mass spectrometer. Subsequently, samples were filtered using TriArt C18 Capillary guard column with 5 um, 5 × 0.5 mm trap column. At the separation step, samples were loaded on TriArt C18 Capillary column, 12 nm, S-3 um, 150 × 0.3 mm for 57 min at a flow rate 5 ul/min. The SWATH acquisition was then performed for 55 min. The time of each scan cycle was 3.2 seconds. The data acquired from each scan cycle (400-1250 m/z) were processed using SCIEX version 1.7.1 software. Three biological replicates for each condition (young vs senescent) were prepared for analysis. For each biological replicate, three technical replicate LC MS runs were undertaken. This results in 9 total replicates for each condition.

### Analysis of the Senescent Proteome

PeacView 2.2 was used to obtain raw counts of peptide distribution which were normalized on peak areas using Marker View 1.2. To calculate the fold change between young vs. senescent cells, a t-test of the nine senescent cell samples against the 9 young samples followed by a 2 sample t-test for each experiment and per gene was carried out. This output contains fold changes as well as p-values. The p-adjusted values were then calculated using the Benjamini-Hochberg procedure in R.

### Functional pathway enrichment-Reactome

Statistical testing for overrepresentation or enrichment of REACTOME terms was performed using the R package Reactome Pathway Analysis version 1.38.0 [21] with the conditions pvalueCutoff = 0.05, pAdjustMethod = “BH”, qvalueCutoff = 0.2, minGSSize = 10 and maxGSSize = 500.

### Immunofluorescence Microscopy

Cells were then fixed in 4 percent paraformaldehyde for 15 min at room temperature prior to permeabilisation in 0.5 pc Triton X-100 for 20 min. Cells were incubated in blocking buffer (3 pc BSA in PBS) for a minimum of 30 min. Primary (SAHF, 1:500; proteintech AB8898) and secondary antibodies (Cy3 conjugated anti-rabbit; Jackson Immunoresearch 711-165-152) were diluted in blocking solution and incubated with the cells for at least 1 hour. Cells were stained with DAPI staining solution for 15 minutes (40 ng/ml DAPI in PBS) and the coverslips were subsequently mounted with Vectashield mounting media (Vector Labs H-1900). Imaging was performed using a Zeiss 880 line scanning confocal microscope.

### RNA Isolation, Sequencing and Analysis

Total RNA was isolated from cultured cells using Trizol reagent (Sigma). Library preparation and sequencing was performed the DBS-Genomics sequencing facility using polyA selection and TrueSEQ library preparation kit (Illumina) and run to obtain 100 bases of paired end reads. The quality of the raw data was controlled with FastQC [22], reads found to be sub-optimal were removed from the analysis pool, for example adapters and bases with an overall quality below 15 in a sliding window of size 4bp were trimmed with Trimmomatic-0.38 [23]. Annotation of the data to the human genome (GRCh38) was carried out using STAR-2.7.0f [24]. Fold changes with corresponding p-adjusted values were calculated using DESeq2 [25].

## Results

### Doxorubicin Induces Senescence in U2OS Cells

WT U2OS cells were harvested on Day 0 (when Doxorubicin was added), Day 2 (when doxorubicin was removed), Day 5, Day 7 and Day 9 prior to staining with SA-ß-gal solution. The blue colouration is indicative of a senescent phenotype with the intensity increasing with time after dox treatment. The first senescent cells were detected at Day 2 and the increased intensity of the stain indicated that most cells were senescent by Day 7 (see figure 1 a). At day 7 the cells showed a significant increase in SA-ß-Gal staining (fold change of 4 compared to day 5). The percentage of SA-ß-Gal positive cells was analysed using ImageJ [26]. By day 9 a decline of 1.25-fold in the SA-ß-Gal positive cells was detected compared to Day 7.

**Figure 1:**
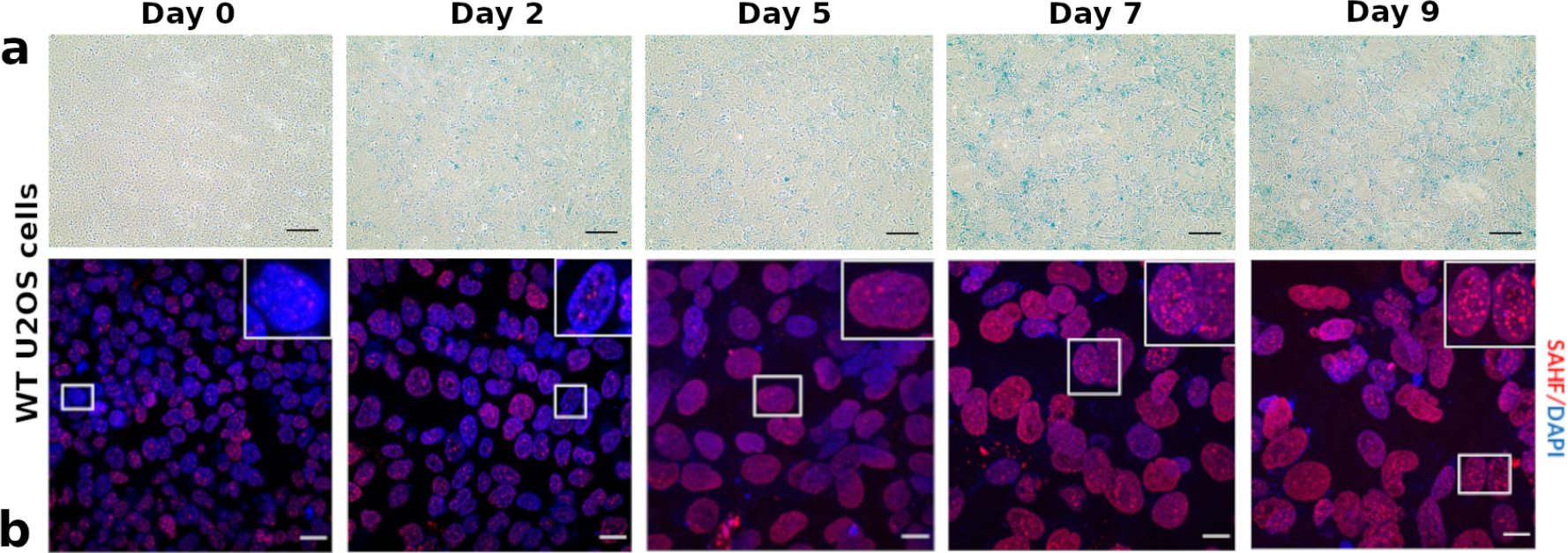
Senescence developed in U2OS cells. a) SA-ß-Gal positive cells were detected in WT U2OS cells. 200 mM Doxorubicin was used to induce cell senescence. Cells were fixed and incubated with SA-ß-Gal staining solution overnight at 37°C. Blue stains were detected in WT U2OS cells at Day 2 post-doxorubicin tatment. Gradual increase in the intensity of blue stain was observed. The blue stain indicates SA-ß-Gal positive (senescent) cells. Scale bar = 100. b) WT U2OS cells were treated with 200 mM doxorubicin for 48 hours. Cells were collected at day 0, 2, 5, 7, 9, fixed and stained with SAHF antibody (red). WT U2OS cells showed no/few SAHF (less than 5 foci per cell) at day 0, 2 and 5. On day 7 and 9, they showed an increased number of SAHF. Scale bar = 20 um.

We employed a second method to assess the senescence induction protocol, by measuring Senescence-associated heterochromatin foci (SAHF). SAHF are formed when the chromatin in the nucleus of senescent cells undergoes remodeling by forming domains of heterochromatin [27]. SAHF formation in U2OS cells was examined following the treatment routine with Doxorubicin. This was followed by immunofluorescence microscopy using a Histone H3 (tri methyl K9) antibody that targets the nucleosome and provides a widely used read-out for SAHF. SAHF formation was monitored on Day 0, 2, 5, 7, and 9 post-doxorubicin treatment in figure 1 b. Cells that are forming 5 or more SAHF are counted and considered senescent. WT U2OS cells showed fewer than 5 structures on Day 0, 2 and 5. On days 7 and 9 the number of SAHF increased and were quantitatively analysed using ImageJ. At Day 7, cells showed a 3.75-fold change increase in the number of SAHF-forming cells of compared to Day 5 cell cultures and a further increase of 2.5-fold change at Day 9 compared to Day 7. Indeed, by Day 9, approximately 47% of U2OS cells were detected to form more than 5 SAHF foci per cell.

### SWATH-MC Proteomics quantitatively identifies Significant Changes in the Senescent Cell Proteome

A total of 5335 proteins were quantified in our measurements, these are listed in table S1 in the supplements. Of these 2672 had a p-adjusted value below 0.05, and are visualized in figure 2 as a volcano plot. Data points in grey represent genes that show less than 2 fold differential expression, i.e. have a | log_2_ *FC*| *<* 1 and are therefore not significantly different between proliferating and senescent cells. Genes with a regulation between 1 *<* | log_2_ *FC*| *<* 2 are significantly expressed and are depicted as small black dots. Highly differentially expressed genes (| log_2_ *FC*| *>* 2) are indicated by large black markers and are also labeled with their gene name. We found 211 up-regulated (log_2_ *FC >* 1) proteins of which 17 were highly up regulated (log_2_ *FC >* 2). Interestingly, only 91 proteins were downregulated (log_2_ *FC <* −1), among these were 6 which were strongly downregulated (log_2_ *FC <* −2). A list of the highly expressed proteins can be found in supplementary table S2.

**Figure 2:**
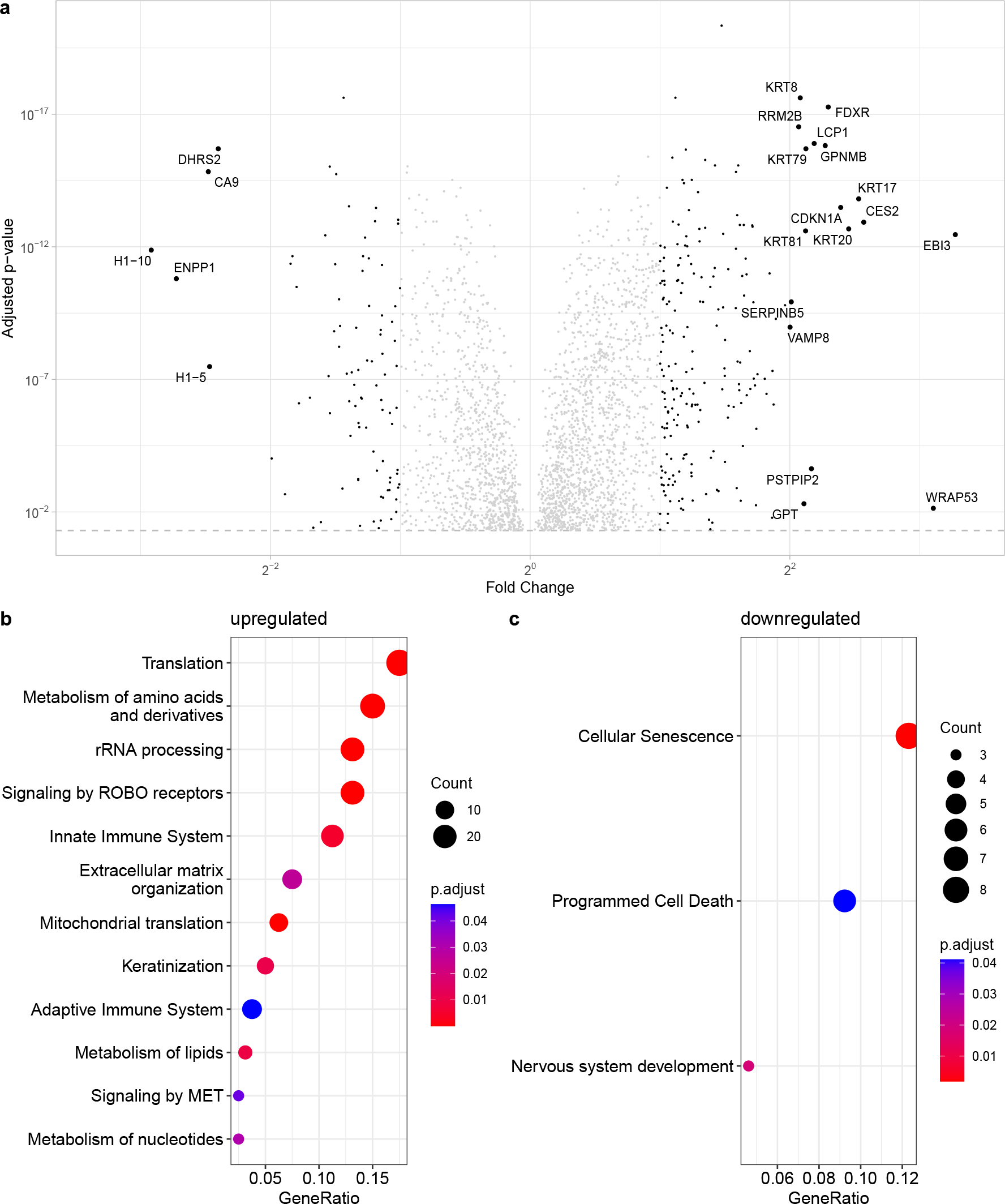
Enrichment analysis of Proteins differentially expressed in young vs senescent U2OS cells. A Volcano-plot for all U2OS proteins with significant p-adjusted value (p-adjusted value ¿ 0.05). Proteins with a |*log*_2_*FC*| *<* 1 are not significant regulated and shown in grey, expressed proteins are significant regulated (1 ¡ |*log*_2_*FC*| ¡ 2) and marked with small black points. Highly expressed proteins (|*log*_2_*FC*| ¿ 2) are shown with big dots and are labeled with the gene name. b and c: Identification of significant over-represented/underrepresented processes in senescence. Figure a shows subsumption of Reactome terms of importance for the upregulated proteins (*log*_2_*FC >* 1), c the significantly underrepresented processes in senescence (underlying proteins have a *log*_2_*FC <* −1).

### The Proteomics Landscape in Senescence

The list of significantly enriched proteins (|log_2_ *FC*| *>* 1, p-adjusted value below 0.05) were investigated with Reactome [28] to uncover functional pathways impacted during senescence. Figure 2b and 2c show the pathways which significantly change in response to senescence induced with doxorubicin. Due to the large numbers of pathway hits, we organised the results such that the parent pathway is emphasised and child pathways that share an enriched parent pathway are ignored. Furthermore, two or more child pathways were merged together in a parent pathway when the parent pathway sufficiently describes the merged child pathways. A list of all pathways can be found in the Supplementary data S3 for the upregulated and S5 for the downregulated proteins. A shortened list where all child pathways are excluded when a parent pathway is enriched can be found in Supplementary S5 (upregulated) and S6 (downregulated), and the list for the figures 2 b and c in supplement S7 and S8, respectively.

As seen in figure 2b, several pathways are enriched in the group of genes upregulated in senescence, such as translation, immune system, extracellular matrix, and metabolism. The numbers of pathways in the downregulated group were more modest and surprisingly included cellular senescence itself. In addition, the downregulated group included apoptosis regulating genes. It has previously been reported that apoptotic pathways are suppressed in senescence.

### Comparison to Existing Senescence Datasets

We next sought to compare the list of senescence related genes from U2OS cells reported here with those previously reported from other cell types. Alvelar et al. developed a comprenhensive database of genes associated with cellular senescence called CellAge. This integrative database utilizes a systems biology approach to the analysis of senescence and has developed gene expression signatures for cellular senescence [29]. We compared our most significantly changed proteins to the CellAge senescence gene expression list. These results are shown in figure 3 and in table S9 in the supplement.

**Figure 3:**
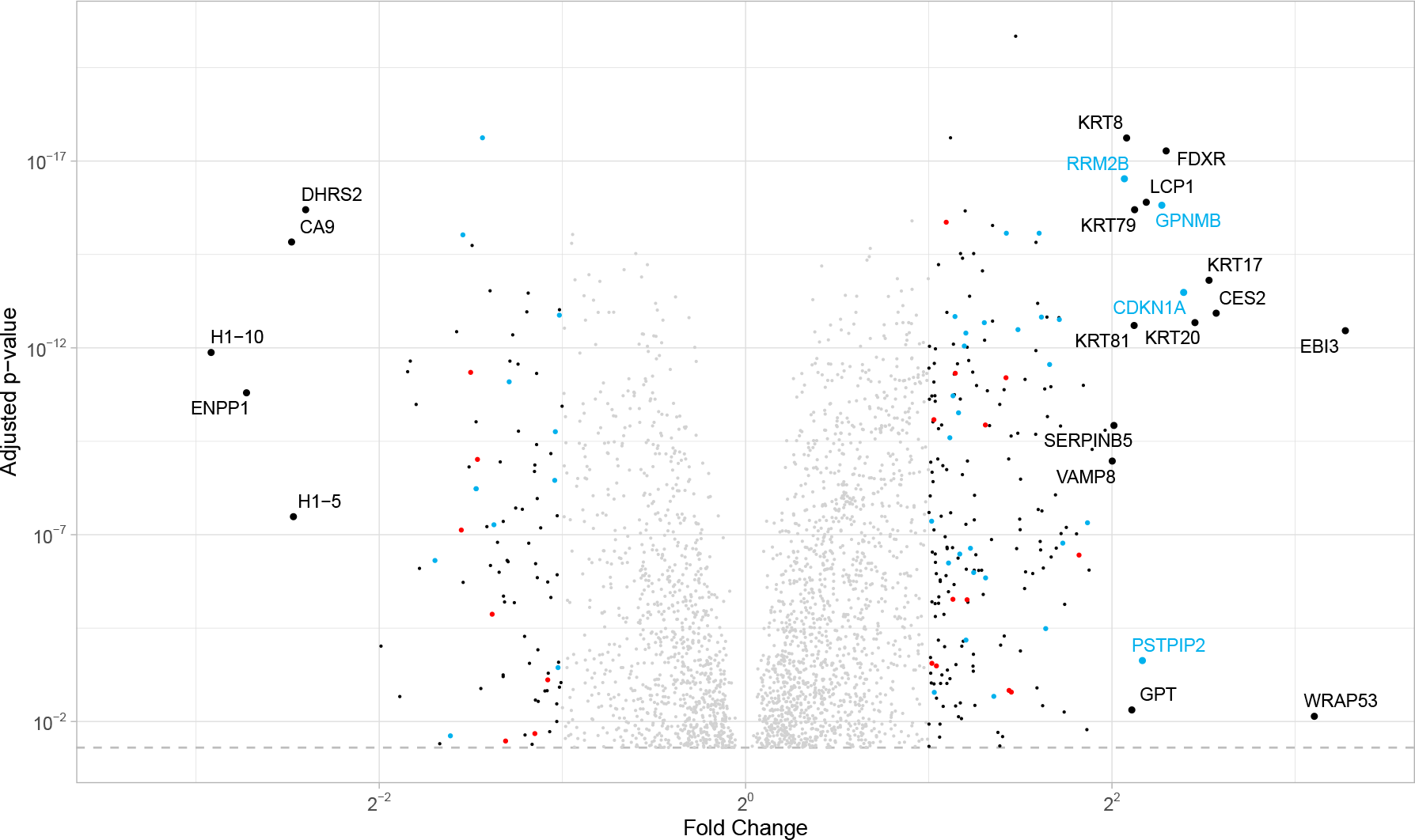
Comparison of the highly regulated proteins in U2OS cells (see figure 2) to the CellAge database. Proteins that have the same direction of regulation (over/under-expressed) in the CellAge database as well as in our data are marked in blue. Proteins with the opposite regulation (i.e. overexpressed in CellAge database but under-expressed in this study) are marked in red. A list with all significant CellAge/HAGR genes in our data can be found in supplementary S9.

Genes that have the same regulation pattern in our U2OS comparison as well as the CellAge list are marked in light blue, while genes that regulate differently (upregulated in one list and downregulated in the other one) are marked in red. Expressed genes (—log2FC— ¿ 1) not included in the CellAge list are marked in black. For highly regulated genes (—log2FC— ¿ 2) the gene name is added. As seen in figure 3 among the genes included in the Cell Age database, especially the highly regulated genes act as expected. The genes that have a contrary regulation mostly have smaller fold changes. We also see several highly regulated genes in our data that are not included in the CellAge list, and are likely cell type specific senescence changes for U2OS cells.

### Large Groups of Genes Undergo Translational Reprogramming in Senescence

The results from quantitative proteomics studies presented thus far were compared with those from RNA-Seq data generated under identical conditions using polyA selected mRNA. Following Illumina sequencing, quality control and further analyses including alignment, gene counts and differential expression, we retained only those genes with a mean count higher than 20 in the young and senescent conditions, fulfilled by 11835 genes. Figure 4 shows a comparison of the fold changes in RNA and protein measurements. In total we had 4927 genes common between RNA and protein datasets. Grey dots depict genes that have a p-adjusted value above 0.05 and therefore do not have enough statistical power to be considered in the analysis. In this group there are 4122 genes. Genes with a p-value below 0.05 which change in both RNA and protein in the same direction (568 genes) are in black. Genes marked in red are statistically significant genes which are upregulated in senescence in RNA but downregulated in the protein measurements (74 genes). Genes marked in blue are regulated the opposite way, downregulated in senescence in RNA but upregulated in the protein measurements (163 genes). Genes with a |log_1_ *FC*| *>* 1 in RNA or in protein of the antiregulated genes are labeled in figure 4 a with gene name. A list with antiregulated genes in either direction can be found in supplementary table S10. Genes with a |log_1_ *FC*| *>* 1 in both conditions, RNA and protein, i.e. highly differentially regulated are few. In the red group there is only one such gene, COMMD8, which is upregulated more than 2 fold at the RNA level but downregulated more than 2 fold in proteomics data. COMMD8 is a potential inhibitor of NF-KB signalling that controls senescence assocoaited inflammatory signalling, hence this disparity in RNA versus proteomics data is interesting. In the blue group there are 9 genes that are highly upregulated at the protein level but more than two fold downregulated in the RNASeq data. These are PHLPP1, ANLN, RACGAP1, KIF23, ITGB4, CCNB1, CAV1, WRAP53 and FAM83D with roles in insulin signalling, cell migration and cell growth.

**Figure 4:**
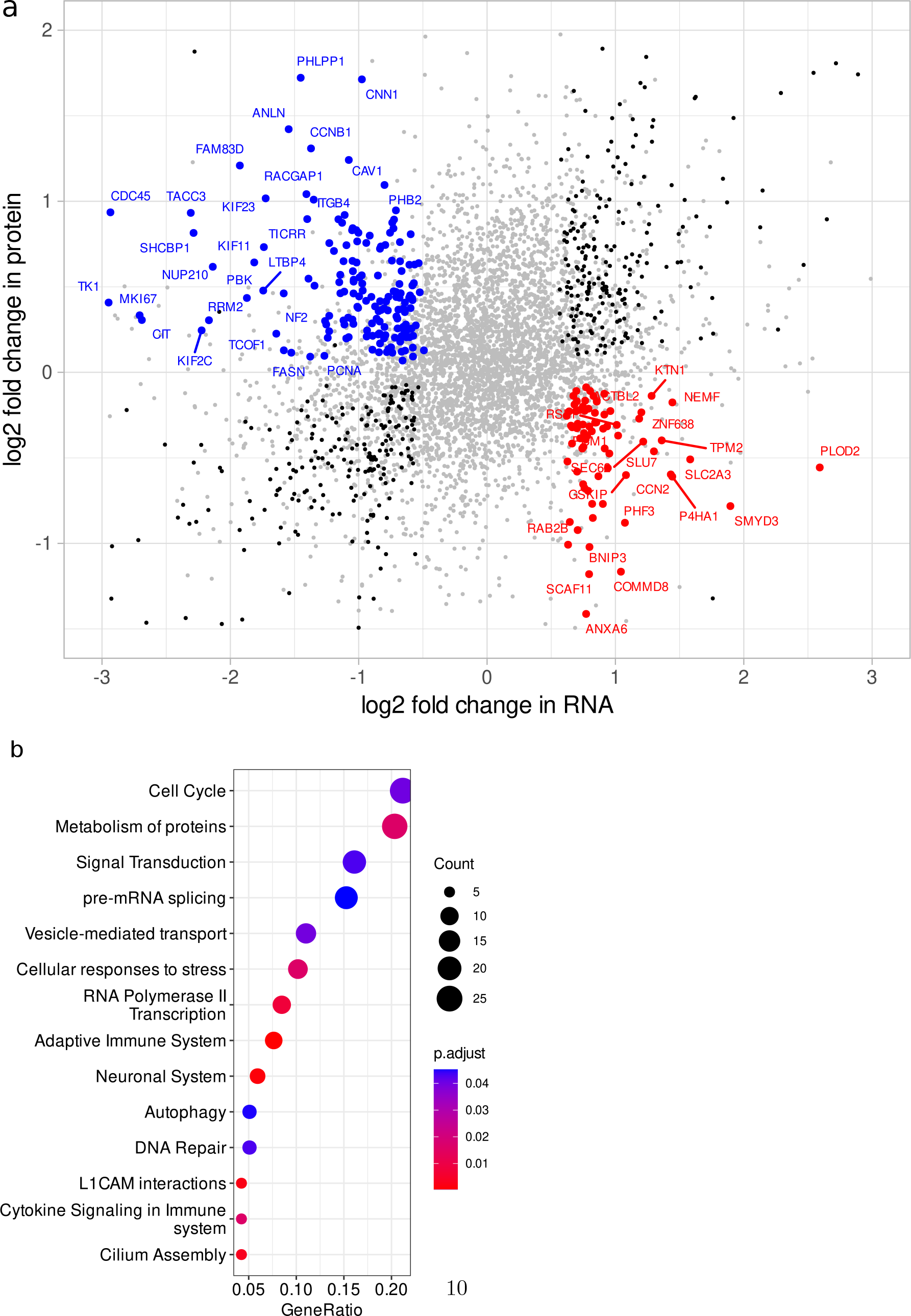
A comparison of the logarithmic fold changes for young vs senescent cells in Protein vs RNA. All genes are included which have a corresponding protein measurement. Grey means that the p-adjusted value was higher 0.05 in RNA or protein. Black genes behave consistent in RNA and protein measurements. Red: A gene is up regulated in the RNA but down regulated in the protein measurements. Blue: Genes which are up regulated in protein but downregulated in RNA. Red and blue genes which have a |*log*_2_*FC*| *>* 1 in at least one condition are labeled. b Reactome pathways which are overexpressed in genes which are upregulated in Proteins and downregulated in RNASeq measurements. There are no enriched pathways for the genes which are downregulated on protein and up regulated10 on RNA level (red genes in a).

A Reactome analysis for the two gene groups with anticorrelated behaviour in 4 was also carried out. No enriched functional pathways were found for the red group in figure 4 a). A summary of the Reactome results for the opposite condition, i.e. genes where RNA levels are downregulated but protein levels are upregulated (blue group in figure 4 a) can be seen in figure 4 b. The pathways and underlying genes can be seen in table S14 in the supplementary material.

## Discussion

### Doxorubicin treatment transforms the proteomic landscape

Cellular senescence is defined by irreversible growth arrest and profound changes in gene expression [2, 30]. Unsurprisingly, proteins involved in senescence and cell cycle progression are amongst the most highly up-regulated proteins in our data set. These include p21, a modulator of cell cycle progression and commonly employed marker of senescence, and RRM2B, a ribonucleotide reductase essential for DNA repair in non-proliferating cells [31] (see figure 2). In addition to the positive SA-ß-gal staining and identification of SAHF, the up-regulation of these proteins confirms the onset of cellular senescence in our model. Among the highly upregulated pathways in the protein data (see figure 2 b), many are connected to SASP and cellular senescence. Furthermore, there are extracellular matrix alterations that are associated with cellular senescence [32]. A further hallmark of cellular senescence is mitochondrial dysfunction which plays important roles not only in the senescence growth arrest but also in the development of SASP and resistance to cell death [33].

Our data also shows that proteins commonly associated with cancer, including prognostic indicators such as SERPINB5, are significantly up-regulated in DNA damage-induced senescence. Examples include WRAP53 which is known to be over-expressed in a variety of cancer cell lines of different origins and promotes cellular transformation [34, 35, 36, 37]. The proteins KRT17 [38], KRT8 [39], KRT20 [40], LCP1 [41], and VAMP8 [42] are also reported to be associated with cancer development/metastasis and cellular proliferation. Functional pathway analysis revealed terms related to translation, inflammation, mitochondrial dysfunction and cell migration to be significantly up-regulated; all of which are typical hallmarks of cancer cells in addition to some being common with senescence. Although considered a *bona fide* supressor of neoplastic transformation, our data suggests that senescence builds a transcriptional/translational landscape that may promote malignancy. Interestingly, we observe a significant enrichment in proteins involved in rRNA processing which is in contrast to previous studies [43]. Ribosome biogenesis and protein translation are finely coordinated and essential for cell growth, proliferation and differentiation. Multiple RP proteins have extra-ribosomal functions including activation of pathways in response to stress, resulting in cell cycle arrest and apoptosis. In cancers, these functions are often misregulated [44]. Several stud-ies have provided evidence for active keratin involvement in cancer cell invasion and metastasis. The kreatinization pathway is here driven by the highly upregulated KRT proteins [45]. Tissue remodeling is promoted by MET tyrosine kinase receptor, which underlies developmental morphogenesis, wound repair, organ homeostasis and cancer metastasis [46]. Several other upregulated pathways are related to cancer. Among these is the metabolism of amino acids and derivatives (GPT) which is here driven by the RPL and RPS gene group and the GTP group of our highly expressed genes. The most significantly down-regulated proteins in our dataset include DHRS2, H1, and ENPP1.

Decreased expression of DHRS2 contributes to p53 stabilization thus promoting the onset of senescence. In mice, Enpp1 has been shown to play a crucial role in regulating aging via Klotho expression and its down-regulation has been shown to be associated with aging [47]. We were initially surprised to find the cellular senescence term to be significantly down-regulated in our data set (see figure 2 c). Among the genes in this pathway is MAP2K6 which is involved in stress-induced cell cycle arrest, transcription activation and apoptosis [48]. Another significant example is ERF which is involved in development, apoptosis, and regulation of telomerase, a key regulator in age-related or replicative senescence. We believe this change may reflect a very late stage of senescence wherein the expression of pro-senescence proteins has reduced and which may represent an incomplete senescent phenotype. To benchmark this study, we compared the proteomic results with known datasets on senescence through the Cell-Age database [29]. Generally, our data is in agreement with that published by Alvelar et al. We believe that any discrepancies observed are likely to be due to differences in cell line and method of senescence-induction, suggesting that this is a useful resource that can support other studies on senescence in this model.

### The Regulation of Translation is Profoundly Altered in Senescence

A large number of pathway terms relating to Translation and rRNA processing appeared amongst the proteins upregulated in senescence in U2OS cells. This led us to compare proteomic data with RNA-Seq which highlights significant irregularities between the detected level of proteins and the expression of their corresponding genes suggesting altered mechanisms of translational regulation between proliferating and senescent cells. Interestingly, our data highlights that genes/proteins where RNA and protein levels are highly anti-regulated (figure 4 a) are typically associated with aging and senescence.

Examples include PHLPP1, which protects against age-related intervertebral disc degeneration [49], and ANLN, the depletion of which induces cellular senescence [50]. Expression of ITGB4 is reportedly down-regulated under oxidative stress or upon inflammatory stimulation leading to the induction of senescence in epithelial cells, mediated through p53 activity [51]. CCNB1 silencing inhibits cell proliferation and promotes cell senescence via activation of the p53 signaling pathway in pancreatic cancer [52], CAV1 which has been shown to induce senescence in resting human diploid fibroblasts [53], and WRAP53 and FAM83D which are commonly over-expressed in a variety of cancers and known to trigger apoptosis [34].

The pathways driven by genes which are up-regulated at the protein level but down-regulated at the RNA level are varied and involve multiple cellular processes. An interesting pathway comprises genes associated with the assembly of the primary cilium, a sensory structure which interprets extracellular signals to stimulate a range of cellular pathways including growth, response to nutrient deprivation, and cellular development [54].

Additional examples include proteins involved in a range of signal cascades and the stimulation of transmembrane receptors. Depending on the cellular context, this may impact cellular proliferation, differentiation and survival. Additional pathways following the same pattern including those related to autophagy, DNA repair and those related to cellular senescence. Cytokines (key components of SASP) and inflammatory mediators are an interesting example of proteins that play a significant role in the senescent phenotype but exhibit striking differences between mRNA and protein levels. These observations may suggest that mechanisms of translational regulation play a more significant role in the induction of the senescent phenotype than transcriptional regulation alone.

## Conclusions

We have employed a cellular model of senescence in a cancer derived cell line to investigate broad transcriptomic and proteomic changes. By combining RNA-seq and proteomic datasets, our data demonstrates a dramatically altered translational landscape at both the protein and RNA level, whilst demonstrating that the model exhibits all the characteristics and markers of the senescent phenotype. The model can be easily implemented and utilised to study senescence in cancer cells in a wide range of contexts. Our data reveals a range of age and disease-relevant proteins and pathways that are altered in senescent cancer cells, such as the assembly of the primary cilium, and highlights the emerging role of lipids in the senescent phenotype. Our data is also suggestive of very significant regulatory changes in translation that warrant further investigation.

## Author contributions

Conceptualization by TS, MA-R, SNG. Data Curation was done by TS and FG. Funding Acquisition, Supervision and Project Administration was undertaken by SNG. Investigation was done by TS and MA-R. Methodology was developed by FG and SNG. Formal Analysis and Software was carried out by FG. Visualization was done by MA-R and FG. The original draft was written by by MA-R and FG and subsequent review & editing by TS and SNG.

## Acknowledgements

We thank Adrian Brown at Durham University, UK, proteomics service for assistance with proteomics. We thank DBS genomics at Durham University, UK, for RNASeq measurements. We thank Nancy Kadesha, Harvard medical school, USA, retired, for sharing the U2OS wild type cell line with us. We thank Anagha S. Setlur and Vidya Niranjan, both RV College of Engineering, Bangalore, India, for helpful comments on the paper.

## Data availability

This project contains the following underlying data:

- RNA data for WT U2OS cells - young and senescent. Available on the European Nucleotide Archive (ENA) under PRJEB59999.
- Proteomics data for WT U2OS cells - young and senescent. Available on Zenodo under the DOI 10.5281/zenodo.7737499.

